# Optimizing Exome Captures in Species with Large Genomes Using Species-specific Repetitive DNA Blocker

**DOI:** 10.1101/2024.04.23.590505

**Authors:** Robert Kesälahti, Timo A. Kumpula, Sandra Cervantes, Sonja T. Kujala, Tiina M. Mattila, Jaakko S. Tyrmi, Alina K. Niskanen, Pasi Rastas, Outi Savolainen, Tanja Pyhäjärvi

## Abstract

Large and highly repetitive genomes are common. However, research interests usually lie within the non-repetitive parts of the genome, as they are more likely functional, and can be used to answer questions related to adaptation, selection, and evolutionary history. Exome capture is a cost-effective method for providing sequencing data from protein-coding parts of the genes. C0t-1 DNA blockers consist of repetitive DNA and are used in exome captures to prevent the hybridization of repetitive DNA sequences to capture baits or bait-bound genomic DNA. Universal blockers target repetitive regions shared by many species, while species-specific c0t-1 DNA is prepared from the DNA of the studied species, thus perfectly matching the repetitive DNA contents of the species. So far the use of species-specific c0t-1 DNA has been limited to a few model species. Here, we evaluated the performance of blocker treatments in exome captures of *Pinus sylvestris*, a widely distributed conifer species with a large (> 20 Gbp) and highly repetitive genome. We compared treatment with a commercial universal blocker to treatments with species-specific c0t-1 (30,000 ng and 60,000 ng). Species-specific c0t-1 captured more unique exons than the initial set of targets, reduced sequencing of tandem repeats, and produced more target regions with high read coverage and narrower depth distribution than the universal blocker. Based on our results, we recommend optimizing exome captures by using at least 60,000 ng species-specific c0t-1 DNA. It is relatively easy and fast to prepare and can also be used with existing bait set designs.

## Introduction

Large and highly repetitive genomes are common among different types of organisms, such as amphibians, fish, and plants. Repetitive DNA sequences can account for close to 80% of the genome in some species, e.g. maize (Flavell et al., 1974). However, research interests usually lie within the non-repetitive parts of the genome, as they are more likely functional and thus can be used to provide answers to questions related to adaptation, breeding, function, and evolutionary history. Non-repetitive parts are also bioinformatically less challenging than repetitive parts. Obtaining sequencing data from regions of interest in a cost-efficient manner is a highly important aspect when working with large genomes. Nowadays various sequencing methods are available. Whole-genome sequencing (WGS) is a great tool for generating large quantities of sequencing data across the genome. However, obtaining enough sequencing depth at specific sites of interest (e.g. genes) for reliable downstream analyses easily becomes expensive and starts to limit the sample size. A lot of sequencing power in WGS is essentially being “wasted” on repetitive DNA sequences, which is not ideal when targeting limited regions of the genome.

Targeted sequencing is a cost-effective alternative to WGS. Restriction-site associated DNA sequencing (RADSeq; Miller et al., 2007) and genotyping by sequencing (GBS; Elshire et al., 2011) methods use restriction enzymes to cut DNA at specific sites. Regions adjacent to these common restriction sites are sequenced. A great advantage of restriction-based methods is that they do not necessarily require any prior knowledge of the studied species. A major downside is that they produce much missing data between individuals due to mutated or missing restriction sites. RNA sequencing is another method that can be utilized in targeted genome analyses, as it limits sequencing to the exons of expressed genes. RNA sequencing data can be also be used to construct a transcriptome (collection of all expressed genes), which can be utilized for other targeted sequencing methods, such as exome capture and amplicon sequencing.

Exome capture (Ng et al., 2009) uses biotinylated baits, designed based on a transcriptome or annotations of a reference genome, to capture and sequence a set of pre-selected exons. Usually, the whole exome can be targeted, as the coding regions only constitute a few percent of the total genomic DNA. Exome captures produce fewer missing data between individuals compared to RNA-sequencing and restriction-based methods.

However, it requires previous knowledge on the studied species to produce a reliable set of baits. Amplicon sequencing is an alternative method to exome capture for sequencing specific genomic regions. This method utilizes baits with polymerase chain reaction (PCR) to produce ultra-deep sequencing of the targeted regions. Target sets are usually smaller than in exome captures. Amplicon sequencing requires less starting DNA material and has simpler DNA preparation steps compared to exome capture. In addition, amplicon sequencing produces higher on-target rates, but lower uniformity compared to exome captures (Samorodnitsky et al., 2015). Even more specific methods, such as primer extension target enrichment (PETE) and duplex sequencing (Kennedy et al., 2014; Schmitt et al., 2012), are available when working with a smaller set of targets. These methods achieve lower error rates than amplicon sequencing and exome capture but are currently too expensive to use on a larger scale.

When using target enrichment methodologies, off-target repetitive DNA can end up being sequenced for two reasons. Firstly, different types of short tandem repeats (microsatellites, minisatellites, and satellites) can have short repetitive sequences that non-specifically bind to the baits blocking the targeted binding to the exons of interest. Secondly, the baits are usually much shorter than the genomic DNA fragments, meaning that a large proportion of the fragment remains in single-stranded form even after the bait has bound to it. Consequently, another genomic DNA fragment containing repetitive DNA sequences can bind to the bait-bound fragment, leading to the sequencing of the whole complex.

Exome captures use c0t-1 DNA to block repetitive DNA sequences from binding to capture baits and bait-bound genomic DNA fragments during the hybridization reaction. C0t-1 DNA is a type of DNA containing a large pool of different types of repetitive DNA sequences of various lengths. It has originally been used to study DNA reassociation kinetics to determine the repetitive DNA contents of genomes within various species, such as maize, pea, and sunflower (Flavell et al., 1974). C0 stands for the initial concentration of DNA and t for the DNA reassociation time in seconds (Britten et al., 1974). The amount of reassociation is measured relative to the original DNA concentration. Repetitive DNA reassociates faster than unique DNA sequences due to the higher probability of finding complementary sequences. Thus repetitive DNA was labeled as low c0t and unique DNA as high c0t.

Nowadays (low) c0t-1 DNA is used in the hybridization reaction of exome capture to reduce non-specific binding, as c0t-1 DNA binds to the repetitive DNA sequences of the genomic DNA. Species-specific c0t-1 DNA is routinely used in the exome captures with model species such as humans and mice. Researchers working with non-model species only have different types of universal blockers available. These universal blockers consist of various highly conserved repetitive DNA sequences and thus do not perfectly reflect the repetitive DNA landscape found within the studied species. The use of universal blockers can potentially lead to increased quantities of non-specific binding in exome captures. Large amounts of non-specific binding lead to a decreased amount of unique reads originating from the target regions. Producing species-specific c0t-1 DNA is a relatively easy and inexpensive alternative to using universal blockers and has been used with good results in amphibians with large genomes (McCartney-Melstad et al., 2016). Despite its potential, species-specific c0t-1 DNA is still not used widely in exome captures of non-model species.

In this study, we evaluated the performance of three different treatments for blocking the binding of repetitive DNA sequences to the capture baits and capture-bound genomic DNA fragments in exome captures of *Pinus sylvestris,* a widely distributed conifer species (Durrant et al., 2016) with a large (> 20 Gbp) and highly repetitive genome. One of the treatments used a commercial universal blocker and two of the treatments used species-specific c0t-1 DNA, prepared using an optimized protocol for *P. sylvestris,* in two different quantities: 30,000 ng and 60,000 ng. We developed a novel bait set for *P. sylvestris* based on earlier transcriptome assembly (Ojeda et al., 2019), and used this set to evaluate the performance of the different treatments. We used only a single genotype in all exome captures to avoid the effects of genetic variation between different individuals. Whole-genome sequencing was conducted to estimate how much and what type of repetitive DNA is being sequenced with no blocking treatment and capture. Our main research questions were: I) Does the use of species-specific c0t-1 DNA improve exome captures by producing more unique reads compared to commercial blockers and II) Does doubling the recommended amount of species-specific c0t-1 DNA produce more unique reads and reduce the capture of repetitive DNA sequences?

## Materials and Methods

### Bait set design

Exome capture baits were designed based on *P. sylvestris* gene-level SuperTranscript assembly (https://www.ncbi.nlm.nih.gov/nuccore/GILP00000000.1; Ojeda et al., 2019). Mosaic SuperTranscripts, multi-copy genes, transcripts originating from organelle genomes, and transcripts containing heterozygous SNPs in haploid megagametophyte samples were excluded from the list of transcripts, as described in Ojeda et al. 2019. Transcripts were also filtered based on the expression levels, requiring the sum of TPM (Transcripts Per Million) counts across all 30 RNA-seq samples (Ojeda et al., 2019) to be above ten (https://figshare.com/articles/dataset/Pinus_sylvestris_assembly_Trinity_guided_gene_level_i nformation/13109492/1). The remaining 108,861 transcripts were considered potential targets for baits. The list of potential targets includes a hand-curated list of 2435 candidate genes for phenology and for primary and secondary metabolism pathways active during heartwood formation (Supplementary File 1).

Based on the list of potential targets, Roche Sequencing Solutions designed a list of candidate baits. Exon-intron boundaries were identified by mapping Roche-designed target areas to the unmasked *Pinus taeda* reference genome v2.01 (https://treegenesdb.org/FTP/Genomes/Pita/v2.01/genome/) (Zimin et al., 2017) using blastn (Altschul et al., 1990). Since *P. sylvestris* does not have a public reference genome, *P. taeda* reference was used in the mapping, as it was the highest quality reference genome available from a related species at the time of bait set design. The mapping process was parallelized using the workflow management software STAPLER (Tyrmi, 2018). Exon-intron boundary was inferred based on alignments where the best alignment did not span the whole target length (https://github.com/tyrmi/PGU/tree/master/find_exon_boundaries). Baits overlapping with any of the 10,748 identified exon-intron boundaries or mapping to highly repetitive regions were excluded from the design. Baits were allowed up to 20 close matches to the reference genome. The final bait set design: SeqCap Design PiSy_UOULU (Supplementary File 2) covers 18,516 regions with a total capture space of 9,345,488 bp.

### DNA extractions and library preparations

A single *P. sylvestris* genotype (E1101), chosen based on the availability of additional sequencing data from the same individual, was used in the exome captures to avoid the effects of genetic variation between different individuals. Before each DNA extraction, 100 mg of frozen needles were cut into < 5 mm pieces and disrupted using TissueLyser (Qiagen) at 30 1/s for 2×120 s. The disrupted samples were incubated in SP1 buffer at 65°C for one hour, followed by DNA extraction using E.Z.N.A SP Plant DNA Kit (Omega Bio-tek). To increase DNA yield, two additional steps were performed: NaOH treatment for the column before loading the sample, and double elution to the same supernatant. DNA concentration was measured using Quant-iT PicoGreen dsDNA kit (Thermo Fisher Scientific).

DNA extraction was followed by DNA fragmentation. Samples were first diluted to 20 ng/µl concentration using TE buffer (1 mM Tris-HCl, pH 8.0, 0.01 mM EDTA). Sample DNA was then fragmented using Bioruptor UCD-200 (Diagenode) for 2×15 min + 13 min in 30/90 s on/off-cycles with power setting "L". DNA fragmentation was followed by double-sided size selection using Agencourt AMPure XP beads (Beckman Coulter) to extract fragments in the desired 250–450 bp range. Beads to sample volume ratio of 0.7 was used for the left side selection and 0.125 for the right side selection. The fragment size distribution was quantified using Bioanalyzer 2100 (Agilent Technologies) with Agilent DNA 12000 Kit (Agilent Technologies).

Libraries were prepared using Kapa HyperPrep Kit (Kapa Biosystems) with SeqCap Adapter Kit (Roche). Ampure XP beads were used in all of the bead-based cleanup steps. DNA concentrations were measured (Quant-iT PicoGreen dsDNA Kit) before the amplification step to determine the minimal number of required PCR cycles. All libraries underwent three cycles of PCR. DNA fragment size distribution for libraries were analyzed using Bioanalyzer 2100 with Agilent High Sensitivity DNA Kit, and DNA concentrations were measured using Quant-iT PicoGreen dsDNA Kit.

### C0t-1 DNA preparation

Species-specific c0t-1 DNA was prepared for *P. sylvestris* according to the protocol published by Zwick et al., (1997) with the following modifications. 1. DNA was fragmented using Bioruptor UCD-200 (1x10 min + 1x6 min, 30/90 s on/off-cycles, power setting “M”) instead of using an autoclave. 2. The targeted fragment size distribution was lowered from the 100– 1000 bp range to the 100-500 bp range (peak between 300-500 bp) to match the size of our library fragments better. Fragmented DNA was diluted to 500-600 ng/µl concentration instead of the recommended 100-500 ng/µl concentration due to the high DNA concentration of our samples and to keep the total reaction volume low. 3. The activity of S1 Nuclease was stopped by adding EDTA (0.5 M, pH 8.0) and incubating at +70°C for 10 minutes instead of the suggested phenol extraction, as it is easier and safer to perform with EDTA. 4.The DNA pellet was washed with 70% ethanol three times after the overnight precipitation, instead of just spinning down the pellet. 5. The DNA pellet was dissolved in 10-20 µl of Tris-HCl (10 mM, pH 8.0) instead of 100-200 µl of TE to minimize the volume of c0t-1 DNA added to the hybridization reaction of exome captures.

The main steps of the complete modified protocol are described here briefly, for the full version see (GitHub). DNA was extracted from fresh *P. sylvestris* needles using E.Z.N.A SP Plant DNA Kit and fragmented using Bioruptor UCD-200. DNA was fragmented to < 500 bp long fragments, confirmed by agarose gel electrophoresis (1% agarose gel, 100V, 30-40 min). Fragmented DNA was diluted to 500-600 ng/µl DNA concentration and 0.3 M NaCl concentration using 5 M NaCl and PCR-grade water. NaCl is used to induce breaks in double-stranded DNA. Diluted DNA was incubated at 95°C for 10 minutes to denature it. Denatured

DNA was allowed to reanneal by incubating at 65°C. Incubation time for the reannealing reaction was calculated using the following formula: C_0_t = 1 = mol/L x T_s_, where C_o_ is nucleotides per liter and time (T) is in seconds (Zwick et al., 1997). Repetitive DNA sequences reanneal faster due to their higher probability of finding complementary strands. The remaining single-stranded DNA molecules (non-repetitive DNA) were digested by adding S1 nuclease (Thermo Fisher Scientific) and incubating at +37°C for 15 minutes. The activity of S1 nuclease was stopped by adding EDTA (0.5 M, pH 8.0) and incubating at +70°C for 10 minutes. Glycogen (Thermo Fisher Scientific), sodium acetate, and 100% ethanol were added to the DNA solution. The DNA solution was stored at -80°C overnight. DNA precipitation was repeated three times on the following day. The cleaned DNA pellet was dissolved in 10 µl Tris-HCl (10 mM, pH 8.0). C0t-1 DNA purity and concentration were measured using Nanodrop (Thermo Fisher Scientific). C0t-1 DNA samples were combined to create pools of 30,000 ng and 60,000 ng.

### Exome captures

Six exome captures were performed with three different treatments and two replicates per treatment for blocking the binding of repetitive DNA sequences to the capture baits and to capture-bound genomic DNA fragments. Two of the treatments were conducted using our in-house *P. sylvestris* c0t-1 DNA in two different amounts: 30,000 ng or 60,000 ng. The third treatment was conducted using the commercial blocker SeqCap EZ Developer Reagent (Roche). The exome captures were performed using a SeqCap EZ HyperCap (Roche) kit with the custom bait set. DNA input for all exome captures was 1000 ng and the incubation time for the hybridization reaction was 20 hours. DNA fragment size distribution for the captured libraries was analyzed using Bioanalyzer 2100 with Agilent High Sensitivity DNA Kit. DNA concentrations were measured using Quant-iT PicoGreen dsDNA Kit. Captured libraries were quantified using LightCycler 480 (Roche) with Kapa Library Quant Kit (Roche) and diluted to 10 nM concentration before sequencing. Sequencing was performed at Biocenter Oulu Sequencing Center using NextSeq 550 System (Illumina) with NextSeq 500/50 Mid Output v2.5 kit (300 cycles).

### Whole-genome sequencing

Whole-genome sequencing was conducted to estimate how much repetitive DNA is being sequenced when no capture and blocking treatment are applied. DNA for WGS was extracted from the haploid megagametophyte tissue from seeds of the same genotype (E1101) that was used in the exome captures. Seed germination was induced by placing seeds onto a moist filter paper on a Petri dish and incubating at room temperature overnight. Germinating seeds were then dissected under a microscope to separate the seed coat, megagametophyte, and embryo from each other. DNA extraction, fragmentation, size selection, and quantification were performed in the same way as described in the section "DNA extractions and library preparations". Library preparation was conducted with NEBNext Library Prep Master Mix Set for Illumina (New England Biolabs) with NEBNext Multiplex Oligos for Illumina (New England Biolabs). Ampure XP beads were used in all of the bead-based cleanup steps. DNA fragment size distribution of the library was analyzed using Bioanalyzer 2100 with Agilent High Sensitivity DNA Kit, and DNA concentration was measured using Quant-iT PicoGreen dsDNA Kit. The library was sequenced at the Institute of Molecular Medicine Finland (FIMM) using HiSeq 2500 System (Illumina).

### Read mapping

De-multiplexing and adapter trimming for the raw exome capture sequencing data were performed by Biocenter Oulu Sequencing Center and for the whole-genome sequencing data by FIMM. Raw read quality for each sample was analyzed using FastQC v0:11.8 (Andrews, 2010) and the results were compared between samples using MultiQC v1.9 (Ewels et al., 2016).

*P. sylvestris* does not have a published reference genome available. *Pinus tabuliformis* is the first conifer reference genome containing chromosome level information (Niu et al., 2022) and it is also a closer relative to *P. sylvestris* than *P. taeda* (Jin et at., 2021) (reference used in the bait set design steps), thus it was used as a reference in all of the following bioinformatic analyses. A masked version of *P. tabuliformis* v1.0 genome (DataDryad) was constructed to increase unique mapping to the genome, as many identical sequences were found between different contigs. To construct a masked version of the reference genome, chromosomes were first split back into contigs. Contigs were then aligned within chromosomes and between unplaced contigs using Minimap2 (Li, 2018a). Then alignments were then chained to longer ones using ChainPaf module of Lep-Anchor (Rastas, 2020). A half of the aligning regions of > 10 kb were masked by masking the region in shorter of the two contigs involved in the alignment. The masked *P. tabuliformis* reference was indexed using bwa (v.0.7.17) (Li & Durbin, 2009) index command, and raw reads were mapped to the indexed reference using bwa-mem algorithm with default parameters. The resulting SAM files were converted to BAM files, sorted, and de-duplicated using SAMtools v1.9 (Danecek et al., 2021). Read group tags were added using Picard tools v.2.21.4 (*Picard Toolkit*, 2019) AddOrReplaceReadGroups command.

The number of raw and mapped sequencing reads varied by millions of reads between the different exome capture samples. To incorporate the difference in the number of mapped reads into the analyses, all samples were randomly subsampled to 45 million mapped reads. Subsampling was done for the de-duplicated BAM files using SAMtools view command. Subsampled BAM files were indexed using SAMtools index command with the option -c (CSI index). Regular BAI index format could not be used, as it only supports individual chromosomes up to 512 Mbp in length.

### Coverage and depth analysis

The bait set is based on *P. sylvestris* transcriptome data. To calculate sequencing coverage (proportion of target region having sequencing reads mapping to it) and depth for the target regions, the *P. sylvestris* transcripts used as target regions for baits were mapped to the masked *P. tabuliformis* reference using bwa-mem algorithm with default parameters and then converted to BAM file using SAMtools. Mapped transcripts were filtered based on mapping quality (MAPQ > 20) using SAMtools view command to reduce the chance of incorrect mapping to 1 in 100. The filtered BAM file was converted to a BED file using Bedtools v2.27.1 (Quinlan & Hall, 2010) bamtobed command. Mapped transcripts were annotated to the *P. tabuliformis* reference genome v1.0 annotation file modified for the masked genome (DataDryad) using Bedtools intersect command. Transcripts were categorized according to whether they were located within a genic or nongenic region to estimate the accuracy of locating *P. sylvestris* transcript target regions within *P. tabuliformis* reference, as genic regions are assumed to be highly similar between the two species due to shared evolutionary history (Jin et al., 2021).

Subsampled exome capture BAM files were converted to BED files using Bedtools. Coverage for the targeted regions in each exome capture BED file was analyzed using Bedtools coverage command. The average depth per target region in each sample was calculated from the output of Bedtools coverage command with option -d. To estimate sequencing coverage variation between different treatments, a random set of 100,000 intervals with the length of 100 bp was generated from masked *P. tabuliformis* reference using Bedtools random command. Coverage for these random intervals was calculated in each sample using Bedtools coverage command.

### Variant calling

A joint variant call for the exome capture subsampled BAM files was performed using BCFtools v1.9 (Danecek et al., 2021) with mpileup and call commands. The masked *P. tabuliformis* genome was used as the reference in the variant calling. Variant calling was used as a tool for comparing the performance of different exome capture treatments. Because only a single genotype was used in the exome captures, all genotype calls are expected to be identical between the different treatments. SNPs originating from the differences between sequences of *P. sylvestris* and *P. tabuliformis* (identified by 1/1-homozygous genotype calls) were removed. SNPs containing the same genotype calls across all treatments were removed, as we were only interested in SNPs having different genotype calls between the treatments.

The remaining SNPs were filtered based on the following criteria: SNP quality (QUAL) > 30, total depth (INFO/DP) > 30, and mapping quality (MQ) > 50. All of the variant filtering steps mentioned above were performed using BCFtools view command. VCFtools v0.1.17 (Danecek et al., 2011) --minDP filter was used to set all genotypes having less than five reads supporting them as missing. The total number of missing genotype calls per sample was calculated using VCFtools --missing-indv command. VCF file containing the filtered SNPs was converted to a BED file using BEDOPS v2.4.41 (Neph et al., 2012) vcf2bed command and annotated to *P. tabuliformis* reference genome v1.0 annotation file using Bedtools intersect command.

### Tandem repeats

Trimmed raw sequencing data from all the exome captures and from the whole-genome sequencing were converted from fastq to fasta format using Seqtk v.1.3-r106 (Li, 2018b) seq command. Tandem repeats were first identified from the WGS data by using Tandem Repeats Finder v.4.09.1-1 (Benson, 1999) with the recommended parameters (matching weight 2, mismatch penalty 5, indel penalty 7, match probability 80, indel probability 10, minimum alignment score 50, and a maximum period size 2000) to identify the most common tandem repeats when no repetitive DNA blocker was used in sequencing. Raw reads from exome captures and WGS data were randomly subsampled to 51 million reads to match the sample with the lowest number of reads (30k_B) using Seqtk sample command with seed (-s) set to 10. Subsampled WGS data served as a baseline when comparing exome capture treatments to each other. Tandem repeats finder with the recommended parameters was used to identify tandem repeats in each subsample separately.

## Results

Sequencing results from exome captures and bait set performance

We generated 53.94 Gbp 150 bp paired-end sequencing data from six exome captures (three different treatments and two replicates of each treatment) using a novel bait set. The number of raw reads per exome capture varied between 52,865,561 and 64,657,074 (Table 1.).

**Table 1:** Exome capture read and mapping statistics.

| Sample | Raw reads | Duplicate reads | Mapped reads |
| --- | --- | --- | --- |
| 60K_A | 58,078,241 | 6,369,885 | 51,435,497 (99.27%) |
| 60K_B | 57,104,428 | 5,772,540 | 51,076,900 (99.30%) |
| 30K_A | 55,188,928 | 7,483,552 | 47,412,572 (99.18%) |
| 30K_B | 52,865,561 | 6,385,977 | 46,571,837 (99.26%) |
| DR_A | 58,235,220 | 8,309,142 | 49,627,974 (99.22%) |
| DR_B | 64,657,074 | 8,118,184 | 56,130,067 (99.10%) |

\* 60K = 60,000 ng species-specific c0t-1 DNA treatment, 30K = 30,000 ng species-specific c0t-1 DNA treatment, and DR = Developer Reagent treatment (universal blocker). A and B are treatment replicate identifiers.

Mapping rates were high, over 99% of reads were mapped in all treatments. The 60,000 ng (60k) c0t-1 DNA treatment had the highest proportion of mapped reads (99.27-99.30%). The number of duplicate reads varied between 5,772,540 and 8,118,184 per treatment (Table 1.), with Developer Reagent (DR) treatment having the largest number of them. One of the 30,000 ng c0t-1 DNA treatment replicates 30k (A) had a poor performance during the hybridization reaction of the exome capture compared to the other c0t-1 treatments, with a noticeably higher number of duplicate reads compared to the other c0t-1 DNA treatments.

By mapping, we successfully identified 18,460 out of 18,516 *P.sylvestris* target regions in the *P. tabuliformis* reference, of which 16,060 were retained after filtering based on mapping quality. Filtered target regions were annotated using the *P. tabuliformis* reference genome annotations. Of the 16,060 filtered target regions, 14,037 (87.4%) were located within annotated genes. In conclusion, target regions were reliably located within the *P. tabuliformis* reference, as they were mapped with high confidence and the majority resided within annotated genes.

### Sequencing coverage and depth in exome captures

The sequencing coverage and depth were analyzed for all target regions in each exome capture treatment. The majority of the target regions were highly covered in all treatments, noticeable as a peak in the coverage distribution near 1.0 (Figure 1A). Another peak in the distribution is located around zero, implying that many target regions did not have any sequencing coverage. The Developer Reagent treatment produced more target regions with low sequencing coverage compared to the c0t-1 DNA treatments (Figure 1B). Both c0t-1 treatments produced more highly covered target regions than the Developer Reagent treatments. However, the overall differences between the treatments for the coverage of the target regions were minor.

**Figure 1:**
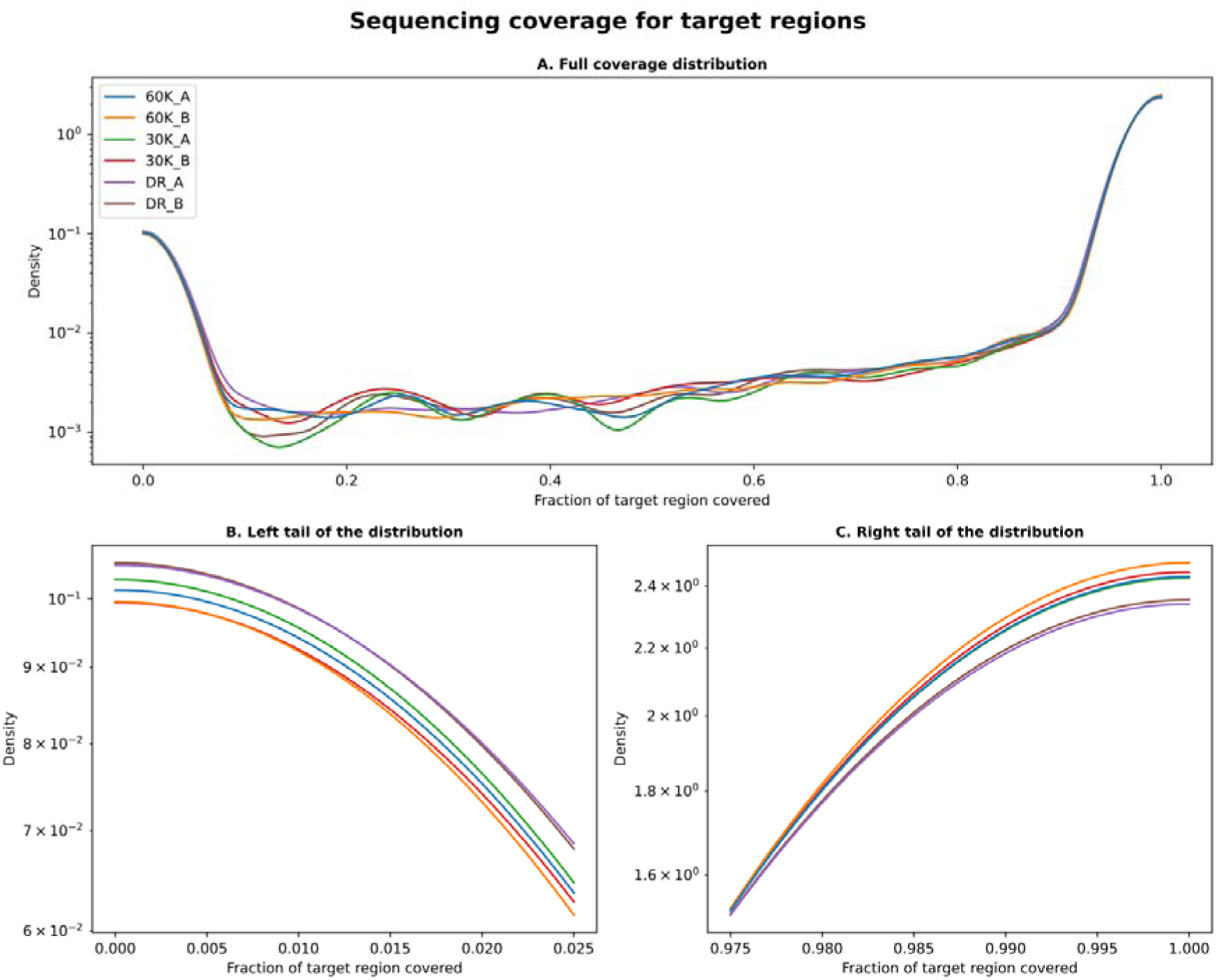
Sequencing coverage for target regions plotted as a kernel density estimate (KDE) plot. A) Full coverage distribution, B) Zoomed view on the left tail of the distribution, and C) Zoomed view on the right tail of the distribution.Y-axis is log scaled.

Sequencing depths within target regions were compared among all treatments (Figure 2). The majority of target region read depths were between zero and 300 for all treatments. The Developer Reagent treatment and 30k c0t-1 DNA treatment replicate A produced target regions with the highest depth values (> 6000) (Figure 2A). They also produced noticeably more depth values ranging from 400 to 900 (Figure 2C). High sequencing depth indicates that much sequencing power has been “wasted” to sequence the same DNA fragments repeatedly, ultimately producing less diverse sequencing libraries. The 60k c0t-1 DNA treatment produced most target regions in the lower end of the depth range (20-200) and had the narrowest depth distribution (Figure 2B). The 30k c0t-1 DNA treatment (B) has a similar depth value distribution as the 60k treatment. The narrower shape of the depth distribution is desired in exome captures as it makes controlling sequencing depth easier, especially with a high number of multiplexed samples.

**Figure 2:**
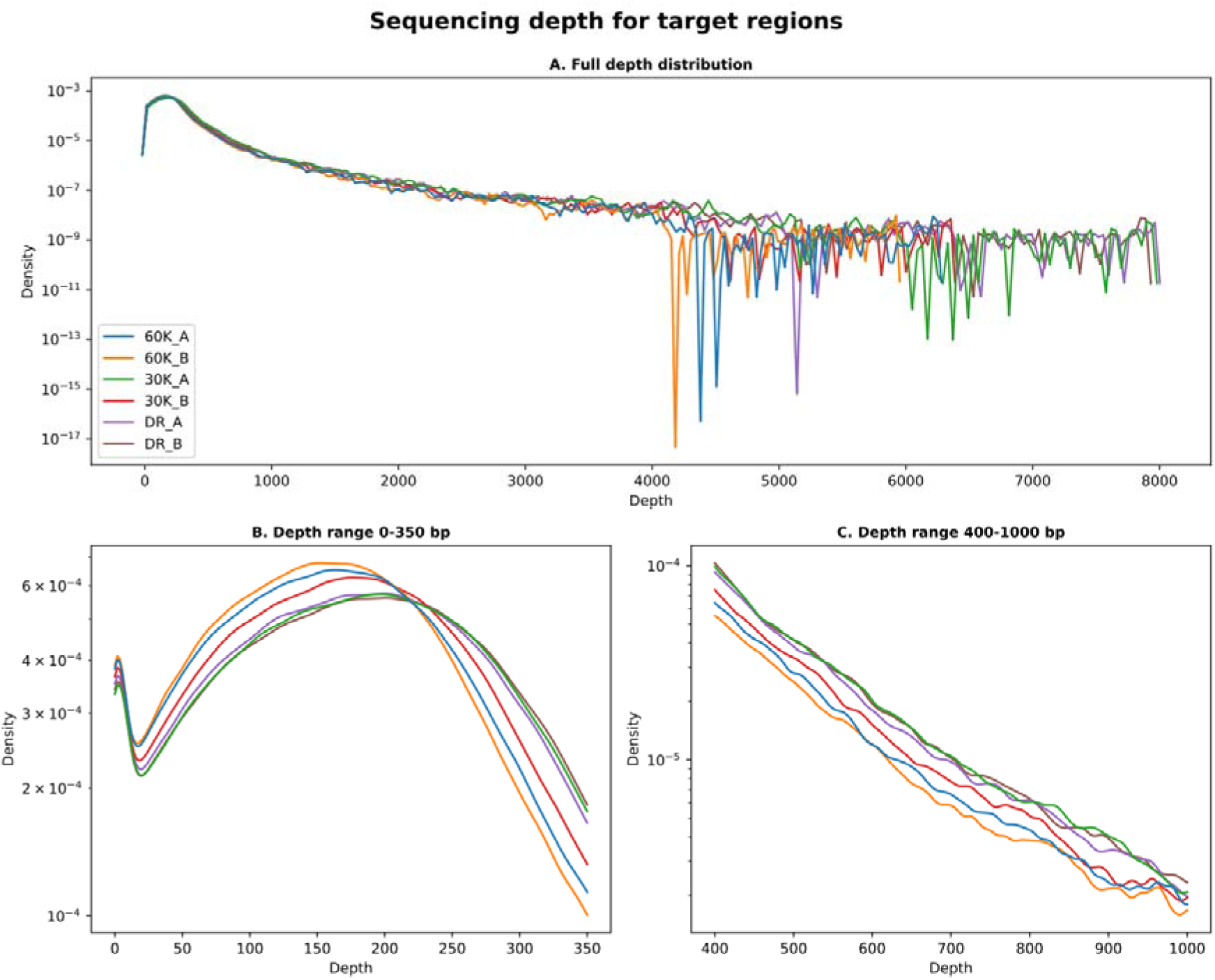
Sequencing depth for target regions plotted as a kernel density estimate (KDE) plot. A) Full depth distribution, B) Zoomed view on the distribution of depth values between 0-350X, and C) Zoomed view on the distribution of depth values between 400-1000X. Y-axis is log scaled.

Sequencing coverage was analyzed in random locations around the *P. tabuliformis* genome to estimate the extent of off-target capture between the different treatments. Coverage was calculated for 100,000 randomly chosen 100 base pair windows. The 60k and 30k (B) c0t-1 DNA treatments produced higher coverage for the random 100 base pair windows compared to the Developer Reagent treatment (Figure 3). These treatments also had fewer windows with low or no sequencing coverage. Analysis of the randomly chosen windows showed that c0t-1 DNA treatments produced more sequencing coverage outside targeted regions than the Developer Reagent treatment. The c0t-1 DNA treatment results were more similar to each other than to the Developer Reagent. Also, their performance did not differ in relation to the amount of c0t-1 DNA used, indicating that the differences in the amount of c0t-1 DNA are not as significant as using c0t-1 DNA instead or Developer Reagent in exome capture.

**Figure 3:**
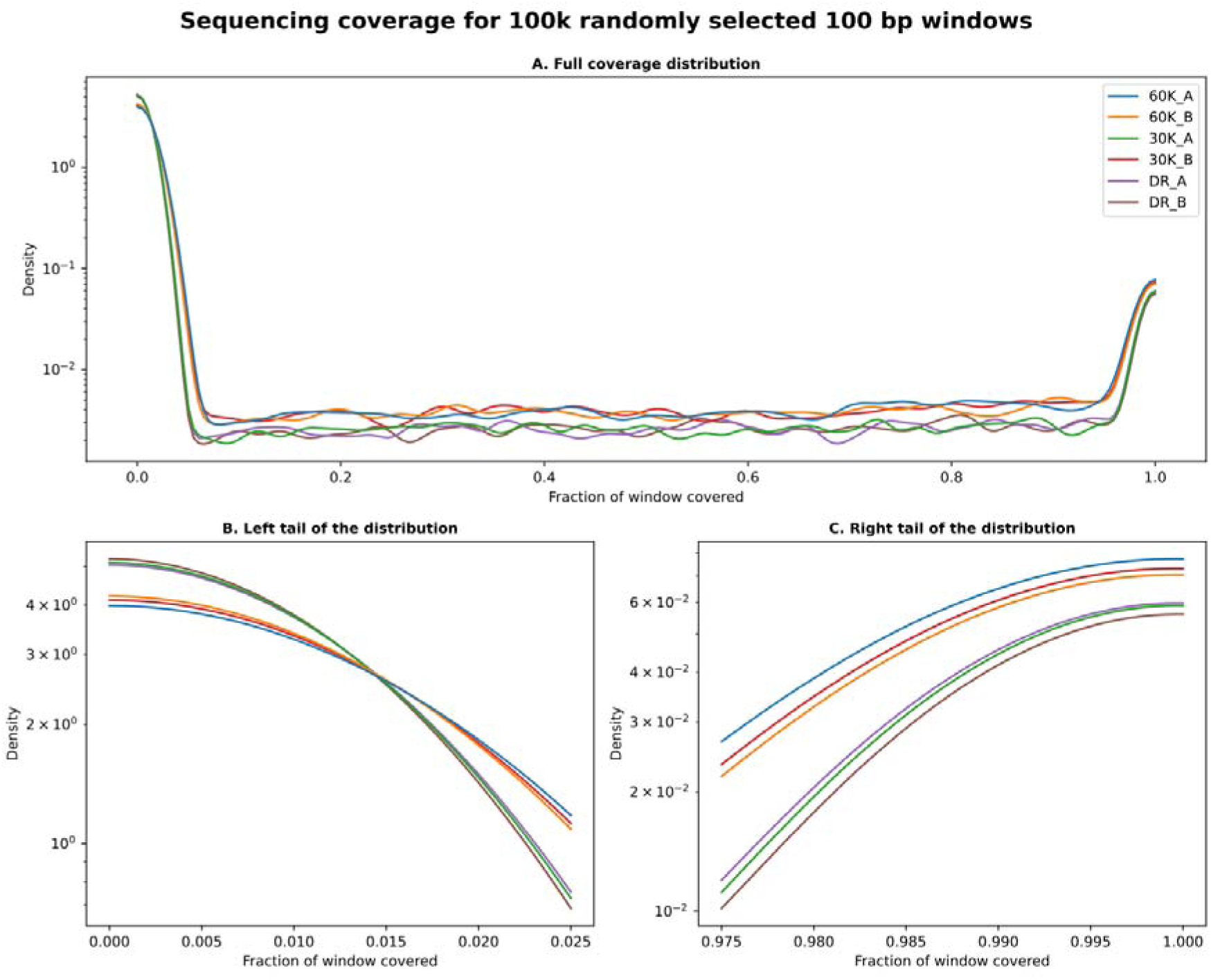
Sequencing coverage for 100,000 randomly chosen 100 bp windows across the genome. A) Full coverage distribution, B) Zoomed view on the left tail of the distribution, and C) Zoomed view on the right tail of the distribution. Y-axis is log scaled.

### Variant call

Variant call was used as a tool for comparing the performance of different exome capture treatments to each other by looking at the genotype calls. All exome captures used DNA from the same genotype, and therefore all genotype calls were expected to be identical across the treatments. A total of 43,955,983 variants were detected (SNPs + indels), of which a set of 535,973 SNPs remained after the filtering steps. Only a small portion of the detected SNPs resided within the target regions (69,278). The rest of the SNPs (466,695) were located outside targeted regions scattered around the *P. tabuliformis* reference genome. Out of these off-target SNPs, 29.3% were located within annotated genes and 44.1% within 10,000 bp of annotated genes, indicating that a significant part of the off-target capture consisted of exons.

The low number of SNPs found in the target regions compared to the off-target regions implies that target regions generally had enough depth in all of the treatments to produce confident genotype calls. This was confirmed by looking at the proportion of missing genotype calls (Figure 4A-B) and genotype depth values (Figure 5A-B) for SNPs located within and outside the target regions. The SNPs within target regions had significantly fewer missing genotype calls compared to the off-target regions. Differences between the treatments are small for the proportions of missing genotype calls for the SNPs within target regions (Figure 4A). However, striking differences between the treatments were observed, when the proportions of missing genotype calls were compared for the SNPs located outside target regions (Figure 4B). The Developer Reagent treatment had about eight times larger proportion of missing genotype calls compared to the 60k c0t-1 DNA treatment. The 30k c0t-1 DNA treatment also had over two times larger proportion of missing genotype calls compared to the 60k treatment. Similar observations can be made when the genotype depth values are compared for the SNPs located outside target regions, as the 60k c0t-1 DNA treatment had the highest median genotype depth for SNPs located outside the target regions compared to both the 30k c0t-1 DNA treatment and Developer Reagent (Figure 5B). Together these results from the variant call indicate that the c0t-1 DNA treatments capture more exons outside the initial set of targets compared to the Developer Reagent treatment. The 60k c0t-1 DNA outperforms the 30k treatment in this aspect. Differences between the treatments are minimal for the SNPs found within the target regions (Figure 5A). However, it is noteworthy that the 60k c0t-1 DNA treatment had fewer missing genotype calls for these SNPs compared to the Developer Reagent treatment while it also had the lowest median genotype depth. Thus, using more c0t-1 DNA reduces excessive sequencing of the same exons while still providing enough depth for confident variant calling.

**Figure 4:**
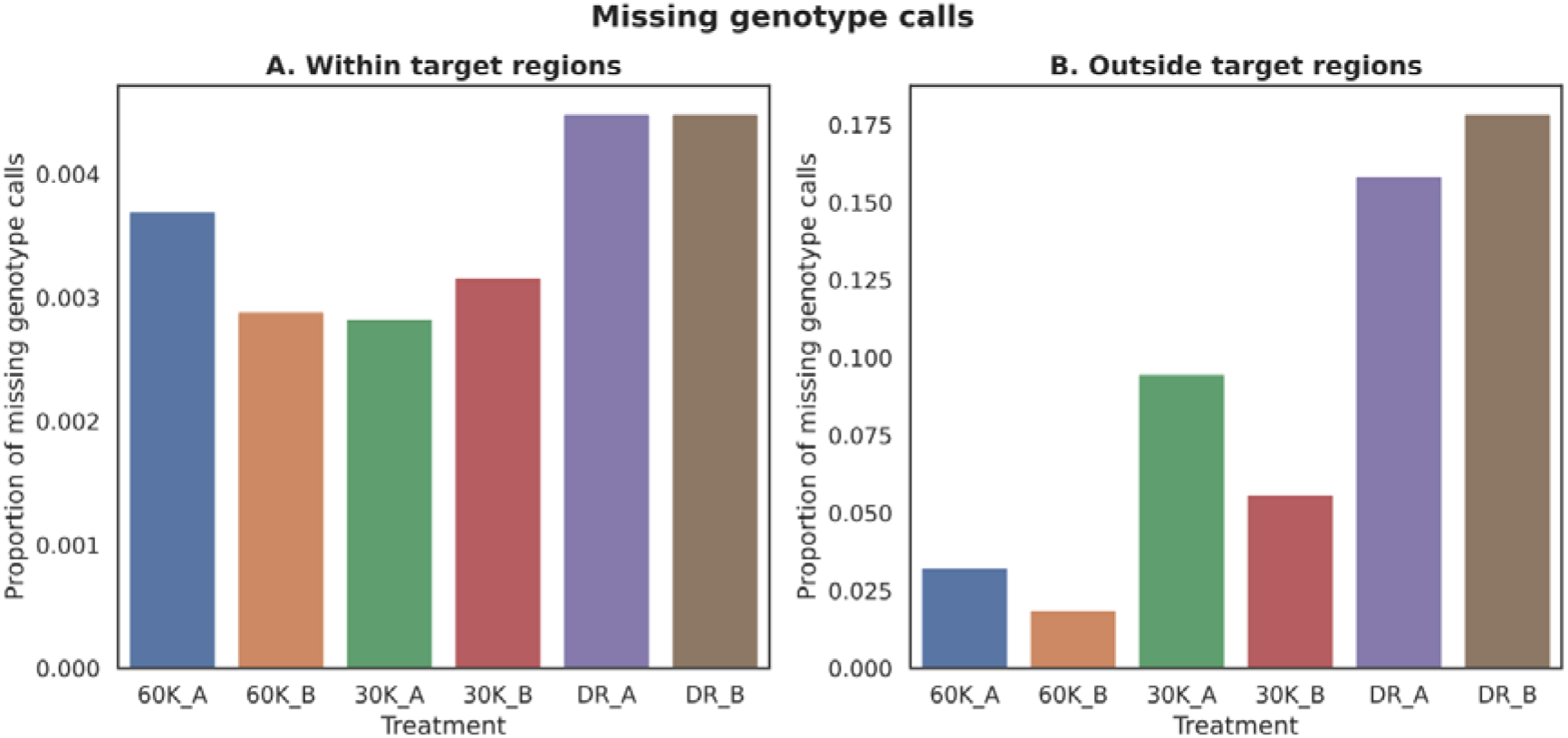
The proportion of missing genotype calls for detected SNPs. A) Within target regions (69,278 SNPs in total), and B) Outside target regions (466,695 SNPs in total). The histogram bars are the treatment and the y-axis gives the number of missing genotype calls divided by the number of SNPs found within the target regions (A) or outside the target regions (B).

**Figure 5:**
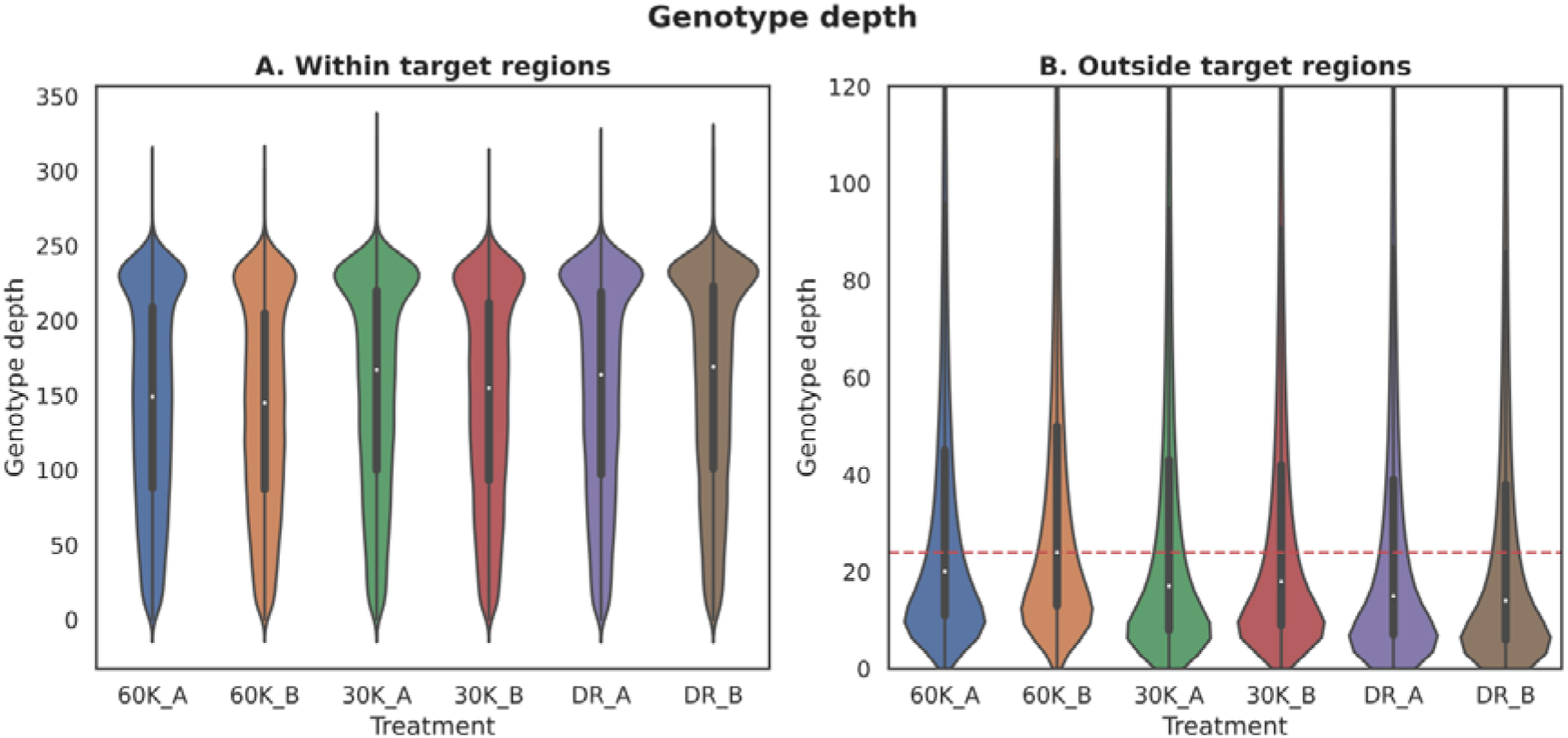
Distribution of genotype depth values for detected SNPs. A) Within target regions (69,278 SNPs), and B) Outside target regions (466,695 SNPs). White dots represent median values for each treatment and black bars represent the values between the first and third quartiles. The red dashed horizontal line is the median depth for the treatment with the highest median genotype depth (60K_A). Unfiltered genotype depth values were used, and genotypes labeled as missing had a read depth of zero.

### Tandem repeats

Whole-genome sequencing data was used to identify the most common tandem repeats in the *P. sylvestris* genome (Supplementary File 3), as no treatment was used to block the sequencing of repetitive DNA. Counts for these most common tandem repeats were compared between the exome capture treatments and WGS data (Figure 6). WGS produced a significantly higher number of the telomere repeat TTTAGGG and its different variants compared to exome captures with different treatments. Similar observations were made for the two centromere repeat variants. Telomeric and centromeric repeats are more commonly found in Developer Reagent than in the c0t-1 DNA treatments. However, the number of telomeric and centromeric repeats is considerably smaller in exome captures than in WGS, showing that sequencing of telomeres and centromeres is not a major concern in exome capture. Exome capture produced more copies of mono- and dinucleotide repeats (A)n, (T)n, (G)n, and (AT)n compared to the WGS (Supplementary File 3). The dinucleotide repeat motif (AT)n has three to four times more copies in exome captures compared to WGS (Figure 6), hinting that this repeat motif is located within or close to genes.

**Figure 6:**
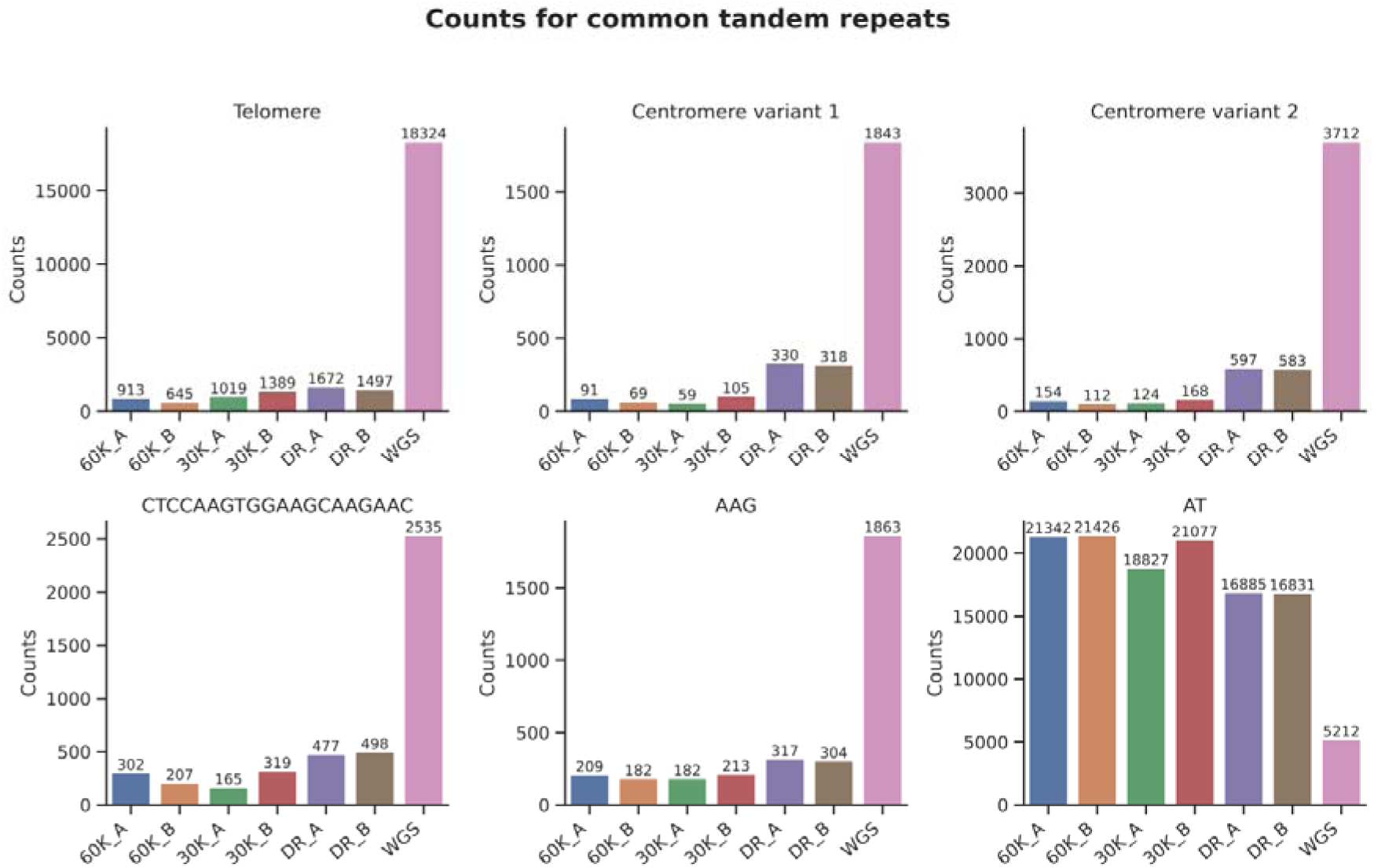
Counts for some of the most common tandem repeats identified in the Whole-genome sequencing data. Telomere repeat contains counts for telomere sequence TTTAGGG and the following variants identified by TRF: TTAGGGT, GTTTAGG, and GGTTTAG. Centromere variant 1 is TGGAAACCCCAAATTTTGGGCGCCGGG and centromere variant 2 is TGGAAACCCCAAATTTTGGGCGCCGCA.

The most common tandem repeats were identified and compared between the different exome capture treatments WGS data. The three most common tandem repeats (longer than 3 bp) in the Developer Reagent treatments had significantly more copies than c0t-1 DNA treatments (Figure 7, upper row). The two most common tandem repeats in the Developer Reagent had over ten times more copies than in the 60k c0t-1 DNA treatment. The third most common tandem repeat had close to three times more copies in the Developer Reagent treatment than the 60k c0t-1 DNA treatment. The Developer Reagent treatment also produced more copies of the most common tandem repeat found in the c0t-1 DNA treatments compared to the 60k c0t-1 DNA treatment (Figure 7, bottom row). The differences between the c0t-1 DNA treatments varied, indicating that the presence of species-specific c0t-1 DNA in exome capture was a more significant factor in the capture and sequencing of tandem repeats than the total amount of c0t-1 DNA used.

**Figure 7:**
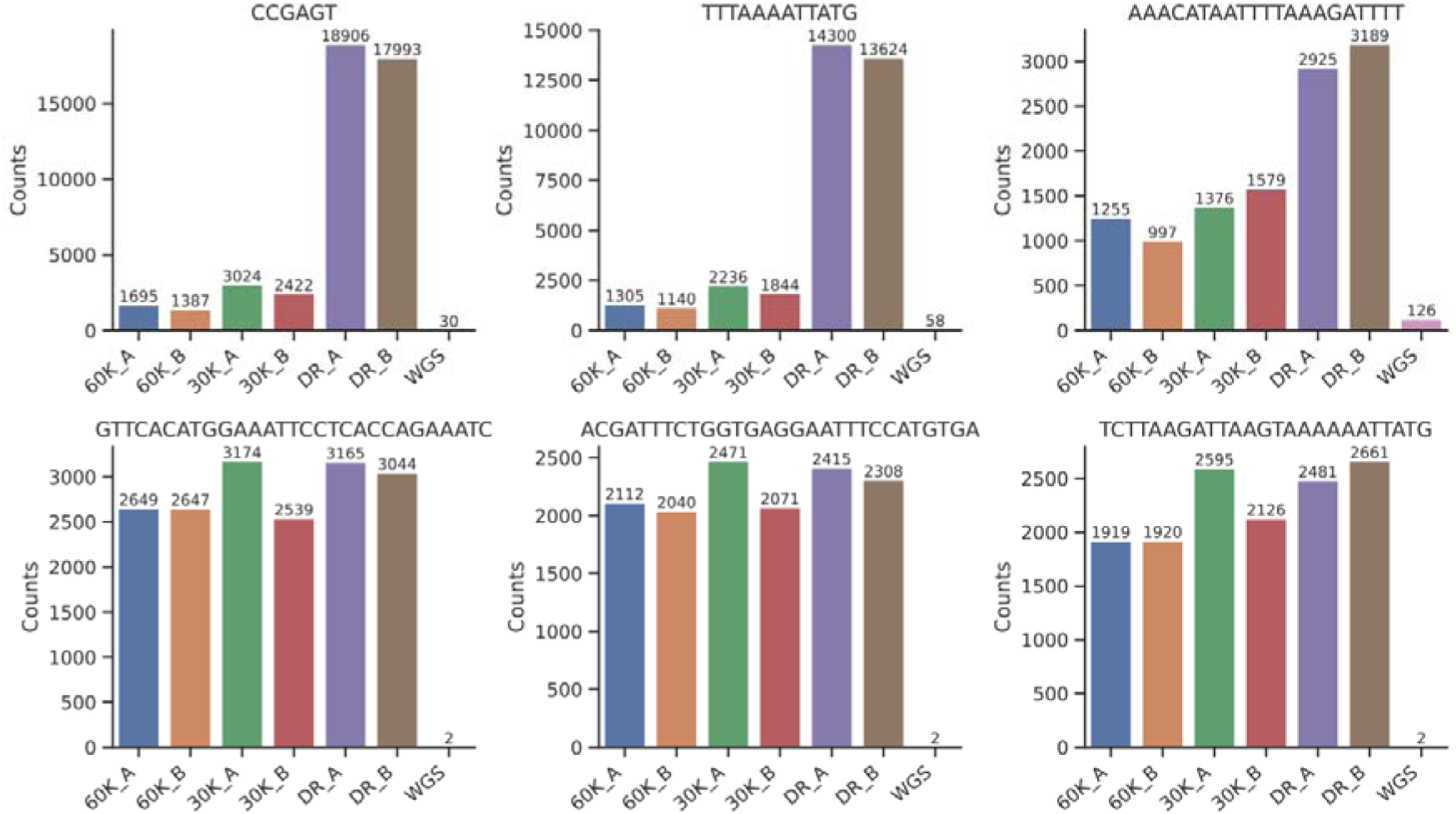
Counts for the most common tandem repeats (longer than 3 bp) found from Developer Reagent treatments and c0t-1 DNA treatments. Tandem repeats in the upper row are the most common tandem repeats found in Developer Reagent treatment and tandem repeats in the lower row are the most common tandem repeats found in both c0t-1 DNA treatments.

## Discussion

The main goal of this study was to compare three different treatments in exome captures for blocking the binding of repetitive DNA sequences to the capture baits and the capture-bound genomic DNA fragments in *Pinus sylvestris*, a species with a large and highly repetitive genome. One of the treatments used a commercially available universal blocker for plants (Developer Reagent) and two of the treatments used species-specific c0t-1 DNA in two different concentrations: 30,000 ng and 60,000 ng. We designed a custom bait set consisting of 18,516 target regions and used it in the exome captures to compare these different treatments to each other. The largest differences between species-specific c0t-1 DNA treatments and the universal blocker treatment were in the amounts of “off-target” capture, where 60,000 ng treatment had by far the best coverage. In the bait set design stage, capture baits were allowed up to 20 close matches to the reference genome (*P. taeda* v2.01), meaning that in reality the captured libraries contained more targets than listed in the bait set design. This is a generally desirable feature, as bait sets are usually not broad enough to target the whole exome. Many of the off-target reads therefore originate from exons, as shown by the large number of off-target SNPs being located within or close to annotated exons. Both species-specific c0t-1 DNA treatments produced higher sequencing coverage in these off-target regions than the universal blocker treatment. The 60,000 ng c0t-1 DNA treatment had the best off-target coverage, shown by its lowest number of missing genotype calls for SNPs located in the off-target areas.

Species-specific c0t-1 DNA treatments also differed from the universal blocker treatment in the amount of captured tandem repeats. Whole-genome sequencing data was used to identify the most common tandem repeats in *P. sylvestris* genome when no treatment was used to block the sequencing of repetitive DNA sequences. The c0t-1 DNA treatments produced fewer copies of some of the most common tandem repeats identified from the whole-genome sequencing data than the universal blocker treatment. The most common tandem repeats found from whole-genome sequencing and the exome captures differed. Exome captures produced more copies of mono- and dinucleotide repeats: (A)n, (T)n, (G)n, and (AT)n compared to the whole-genome sequencing. The high copy numbers of (AT)n repeat motif in exome captures could be partially explained by the sequencing of TATA boxes located close to transcription start sites. The mononucleotide repeat (A)n and many other short tandem repeats have been linked to *cis*-regulatory elements in gene expression regulation in eukaryotes (Horton et al., 2023; Pholtaisong et al., 2022), explaining their abundance in exome captures. However, the abundance of poly-G repeats is a common artifact in two-channel Illumina sequencing systems observed when the dark base G is called after the termination of synthesis. Exome captures underwent two rounds of PCR, which can lead to higher copy numbers of poly-G repeats compared to WGS. The universal blocker treatment produced more copies for the most common tandem repeats found in the exome captures than both c0t-1 DNA treatments. Interestingly, many of the most common tandem repeats found in the universal blocker treatment were found in multiples of three, suggesting they could be potential amino acid repeats. The high number of sequenced amino acid repeats can be a sign of a less diverse sequencing library, which would be in line with our findings from sequencing coverage and variant call analyses.

Species-specific c0t-1 DNA treatments and the universal blocker treatment had little differences in the coverage and read depth of target regions. The bait set had excellent performance, as most of the target regions were completely or close to completely covered by sequencing reads with high sequencing depth in all of the treatments. Only a tiny portion of target regions had low or no sequencing coverage. This was likely caused by incorrect mapping of some *P. sylvestris* target regions to the masked *P. tabuliformis* reference genome. A majority of the genes within the *P. tabuliformis* (91.2%) genome were identified as being duplicated through dispersed duplication (Niu et al., 2022). Mapping reads and target regions to a different duplicate of the same gene can result in some target regions appearing to lack coverage. Both c0t-1 DNA treatments produced more highly covered target regions and fewer target regions with low or no coverage compared to the universal blocker treatment. Both c0t-1 DNA treatments also had a narrower depth distribution with fewer high-depth values (> 400). With multiplexed libraries, a narrower sequencing depth distribution makes controlling per sample sequencing depth easier and leads to reduced over-sequencing of the target regions. Together these results indicate that the species-specific c0t-1 DNA blocks more efficiently the binding and capture of various repetitive DNA sequences, leading to more diverse sequencing libraries with fewer copies of tandem repeats and reduced over-sequencing of target regions.

Based on our results, we recommend using species-specific c0t-1 DNA in exome captures. C0t-1 DNA can be prepared with a short two-day protocol relatively cheaply, as it does not require highly specialized equipment or expensive reagents. We based the lower 30,000 ng of c0t-1 DNA treatment on the results by McCartney-Melstad et al. (2016), who suggested using at least 30,000 ng of c0t-1 DNA per 1000 ng of input DNA. They found that increasing the amount of c0t-1 DNA increased the percentage of unique reads mapping to target regions in amphibian exome captures (McCartney-Melstad et al., 2016). We recommend using at least 60,000 ng of species-specific c0t-1 DNA per 1000 ng of input DNA library in exome captures, as doubling the amount of species-specific c0t-1 DNA used in the exome captures produced more reads from off-target exons and a narrower sequencing depth distribution for target regions with no negative side effects. We speculate that even higher quantities could be beneficial. However, more testing with higher quantities of c0t-1 DNA and different multiplexing strategies would be required. Successful hybridization and capture reactions play an essential role in exome captures, as one replicate of 30,000 ng c0t-1 DNA treatment with subpar performance in the hybridization reaction had contrasting results compared to the other replicate of the same treatment, and the 60,000 ng c0t-1 DNA treatment.

C0t-1 DNA has some interesting applications outside of its use in blocking the binding of repetitive DNA sequences in exome captures. It can be used in fluorescence in situ hybridization (FISH) to locate repetitive DNA sequences within chromosomes (Chang et al., 2008; Sevilleno et al., 2020) or it can be sequenced to analyze the repetitive DNA (low c0t DNA) contents within a genome. This approach has been used to analyze repetitive DNA contents within chicken (Wicker et al., 2005), banana (Hřibová et al., 2008), and beet (Zakrzewski et al., 2010) genomes. Fragment sizes of c0t-1 DNA can be easily adjusted to match the fragment sizes required by the selected sequencing method. Different reannealing times can be used to either only target the highly repetitive DNA sequences or also some moderately repetitive DNA sequences. Repetitive sequence motifs within tandem repeats and interspersed repeats are typically rather short, meaning that regular short-read sequencing methods, such as Illumina sequencing, can be used.

An even more interesting application is the use of high c0t DNA (low copy number DNA) in whole-genome sequencing. Different DNA re-association times can be used to separate repetitive DNA sequences from unique DNA sequences (genes). After certain reassociation time repetitive DNA sequences are in a double-stranded form and unique DNA sequences are in a single-stranded form. The single-stranded DNA can then be separated from the double-stranded DNA by using hydroxyapatite chromatography (HAP chromatography) (Peterson et al., 2002) or by using duplex-specific nuclease to hydrolyze double-stranded DNA (Shagina et al., 2010). Peterson et. al (2002) suggested using HAP chromatography to separate single-stranded DNA from double-stranded DNA after renaturation to specific c0t values to efficiently capture unique sequences from eukaryote genomes. A method based on the same idea of using DNA renaturation to normalize repetitive DNA was developed and successfully tested with maize (Yuan et al., 2003). Shagina et al. (2010) suggested using duplex-specific nuclease extracted from Kamchatka crab to eliminate double-stranded low c0t DNA, as an alternative to using HAP chromatography. Both of these methods offer powerful ways for eliminating repetitive DNA sequences from whole-genome sequencing data sets and are a great alternative to currently popular targeted sequencing methods. As opposed to exome capture, they do not require any prior knowledge about the genes nor are as random as restriction-based methods. These methods appear to have been largely forgotten during the rapid advances in sequencing technologies within the 2000s and should now be revisited.

## Conclusions

We demonstrated that exome capture can be further optimized for species with large and highly repetitive genomes by using species-specific c0t-1 DNA to block the binding of repetitive DNA sequences to the capture baits and to the capture bound genomic DNA fragments during the hybridization reaction. This approach can be used with all existing bait set designs and, furthermore, c0t-1 DNA can be prepared with a short two-day protocol without expensive reagents. Based on our results, we recommend using at least 60,000 ng of species-specific c0t-1 DNA for 1000 ng of input DNA in exome captures, as larger quantities of it were observed to produce more unique reads from exons, especially in the “off-target” regions.

## Acknowledgments

We thank Soile Alatalo for her help and expertise during all the wet laboratory work steps. This work was supported by the following grants: Academy of Finland (287431 and 319313 to Tanja Pyhäjärvi, 343656 to Pasi Rastas, and 309978 to Sonja T. Kujala), Procogen Horizon (289841), Emil Aaltonen Foundation (Nuoren tutkijan apuraha to Robert Kesälahti). The authors wish to acknowledge CSC - IT Center for Science, Finland, for computational resources.

## Data Accessibility and Benefit-Sharing Section

Raw sequence reads are deposited in the SRA (BioProject XXX). The masked *P.tabuliformis* reference genome, annotations with updated genomic coordinates and additional information related to bait set design are available on DataDryad (XXXX).All scripts used in the bioinformatic analyses and the modified c0t-1 DNA protocol are available on GitHub (XXXXX).

## Author Contributions

Tanja Pyhäjärvi, Outi Savolainen, Sonja T. Kujala, Jaakko S. Tyrmi, and Tiina M. Mattila designed the bait set. Timo A. Kumpula and Robert Kesälahti optimized the protocol for producing species-specific c0t-1 DNA for *P. sylvestris*. Robert Kesälahti, Sandra Cervantes, and Tanja Pyhäjärvi designed the experiment. Robert Kesälahti, Alina K. Niskanen, and Tanja Pyhäjärvi analyzed the data. Pasi Rastas created the masked version of *P. tabuliformis* reference genome. Robert Kesälahti wrote the manuscript with the help of Alina K. Niskanen and Tanja Pyhäjärvi.

## Supplementary Data

Supplementary file S1 (list_of_2435_candidate_genes.txt): transcript IDs (Ojeda et al., 2019) for the 2435 candidate genes for phenology and for primary and secondary metabolism pathways active during heartwood formation.

Supplementary file S2 (bait_set_SeqCap Design PiSy_UOULU_target_regions.fasta): Fasta sequences for the 18,516 targeted *P. sylvestris* regions.

Supplementary file S3 (tandem_repeat_counts.xlxs). Counts for the identified tandem repeats from all treatments and WGS data.

